# Modified Vaccinia Ankara Based SARS-CoV-2 Vaccine Expressing Full-Length Spike Induces Strong Neutralizing Antibody Response

**DOI:** 10.1101/2020.06.27.175166

**Authors:** Nanda Kishore Routhu, Sailaja Gangadhara, Narayanaiah Cheedarla, Ayalnesh Shiferaw, Sheikh Abdul Rahman, Anusmita Sahoo, Pei-Yong Shi, Vineet D. Menachery, Katharine Floyd, Stephanie Fischinger, Caroline Atyeo, Galit Alter, Mehul S. Suthar, Rama Rao Amara

**Affiliations:** Emory Vaccine Center, Yerkes National Primate Research Center, Emory University, Atlanta, Georgia 30329, USA; Department of Microbiology and Immunology, Emory School of Medicine, Emory University, Atlanta, Georgia 30322, USA; Department of Biochemistry and Molecular Biology, The University of Texas Medical Branch, Galveston, TX, USA; Department of Microbiology and Immunology, The University of Texas Medical Branch, Galveston, TX, USA; Ragon Institute of MGH, MIT and Harvard, Cambridge, Massachusetts, USA; Department of Pediatrics, Division of Infectious Diseases, Emory University School of Medicine, Atlanta, GA 30322, USA

## Abstract

There is a great need for the development of vaccines for preventing SARS-CoV-2 infection and mitigating the COVID-19 pandemic. Here, we developed two modified vaccinia Ankara (MVA) based vaccines which express either a membrane anchored full-length spike protein (MVA/S) stabilized in a prefusion state or the S1 region of the spike (MVA/S1) which forms trimers and is secreted. Both immunogens contained the receptor-binding domain (RBD) which is a known target of antibody-mediated neutralization. Following immunizations with MVA/S or MVA/S1, both spike protein recombinants induced strong IgG antibodies to purified full-length SARS-CoV-2 spike protein. The MVA/S induced a robust antibody response to purified RBD, S1 and S2 whereas MVA/S1 induced an antibody response to the S1 region outside of the RBD region. Both vaccines induced an antibody response in the lung and that was associated with induction of bronchus-associated lymphoid tissue. MVA/S but not MVA/S1 vaccinated mice generated robust neutralizing antibody responses against SARS-CoV-2 that strongly correlated with RBD antibody binding titers. Mechanistically, S1 binding to ACE-2 was strong but reduced following prolonged pre-incubation at room temperature suggesting confirmation changes in RBD with time. These results demonstrate MVA/S is a potential vaccine candidate against SARS-CoV-2 infection.

## Introduction

The novel SARS Coronavirus-2 (SARS-CoV-2) infection has recently emerged as a pandemic across the world. As of June 24, 2020, the SARS-CoV-2 has infected more than 9.3 million people and over 479,000 people have succumbed to COVID-19, a disease caused by SARS-CoV-2. Thus, there is an urgent need for the development of a vaccine that can rapidly induce anti-viral immunity and prevent infection. Previous data from other related coronavirus infections such as SARS-CoV and MERS-CoV demonstrated that a strong neutralizing antibody response against the spike protein can effectively prevent infection *(1–4)*. Consistent with this, recent studies used DNA, mRNA and chimp adenovirus-based vaccines expressing full length spike as an immunogen and showed induction of neutralizing antibodies against SARS-CoV-2 *(5–9)*. Encouragingly, some of these studies showed protection from disease either in small animals or in rhesus macaques. Emerging data from COVID-19 patients and animal studies show a strong association between antibody response targeted to receptor binding domain (RBD) within the S1 region of the S protein and neutralizing activity suggesting that COVID-19 vaccines should aim to target antibody response to RBD *(10)*.

Modified vaccinia Ankara (MVA) is a highly attenuated strain of vaccinia virus. The safety, immunogenicity, and protective capacity of the replication-deficient MVA has been well-established and has been widely used for developing vaccines against infectious diseases and cancer in preclinical research and humans *(11, 12)*. There are several advantages to MVA-based vaccines. 1) They are safe and well-tolerated including in HIV infected individuals *(13)*. 2) They induce robust antibody responses after a single vaccination and can be boosted at least 10-fold with a second dose *(14–18)*. 3) MVA vaccine-induced antibody responses in humans are durable with little contraction over a 6 month timeframe *(16)*. 4) MVA can be delivered through multiple routes and can be used to generate a mucosal antibody response. 5) MVA can accommodate large inserts (>10kb) that will allow expression of multiple antigens in a single vector. 6) MVA vectored recombinants are stable and can be produced at high titer enabling ease of vaccine manufacturing. 7) MVA vaccines can induce CD4 and CD8 T cell responses important for protection against some viral infections *(14)*. 8) MVA-based vaccines have been shown to protect against SARS-CoV, MERS-CoV, Zika virus and Ebola virus *(4, 17–19)*.

In this study, we developed two modified vaccinia Ankara (MVA) based vaccines which express either a membrane anchored full-length spike protein (MVA/S) stabilized in a prefusion state or the S1 region of the spike (MVA/S1) which forms trimers and is secreted. Both immunogens contained the receptor-binding domain (RBD) which is a known target of antibody-mediated neutralization. The MVA/S also incorporated two mutations that have been shown to maintain the spike protein in a prefusion confirmation *(20, 21)*. Using a mouse model, we evaluated the antibody responses against the SARS-CoV-2 spike protein following immunization with these two MVA-based vaccines.

## Results

### The MVA vaccines express high levels of full-length stabilized spike and trimeric soluble S1 proteins

To develop the MVA recombinants we synthesized the full-length spike gene (amino acids 1-1273) with stabilizing mutations (K986P, V987P) or the S1 region with a small portion of S2 region (amino acids 14 to 780). To promote active secretion of the S1, we replaced the first 14 amino acids of the spike sequence with the signal sequence from GM-CSF (**Fig. 1A**). Both sequences were optimized for MVA codon usage, corrected for poxvirus transcription termination sequences and cloned into pLW73 vector that will allow us to insert the recombinant sequences under mH5 promoter in the essential region of MVA. The recombinants were selected as described previously and characterized for protein expression by flow cytometry and Western blotting. As expected, the MVA/S expressed high levels of spike on the cell surface (**Fig. 1B**) and the expressed protein had a molecular mass of about 180 kDa (**Fig. 1C**). Similarly, the MVA/S1 was expressed intracellularly (**Fig. 1B**), and a molecular mass of about 114 kDA was also secreted into the supernatants (**Fig. 1C**). The spike protein expressed by MVA/S on the surface seemed folded correctly based on strong binding to ACE2 (**Fig. 1D**). Interestingly, the S1 protein was found to form trimers based on gel filtration profile and native-PAGE analysis (**Fig. 1E**).

**Fig 1.**
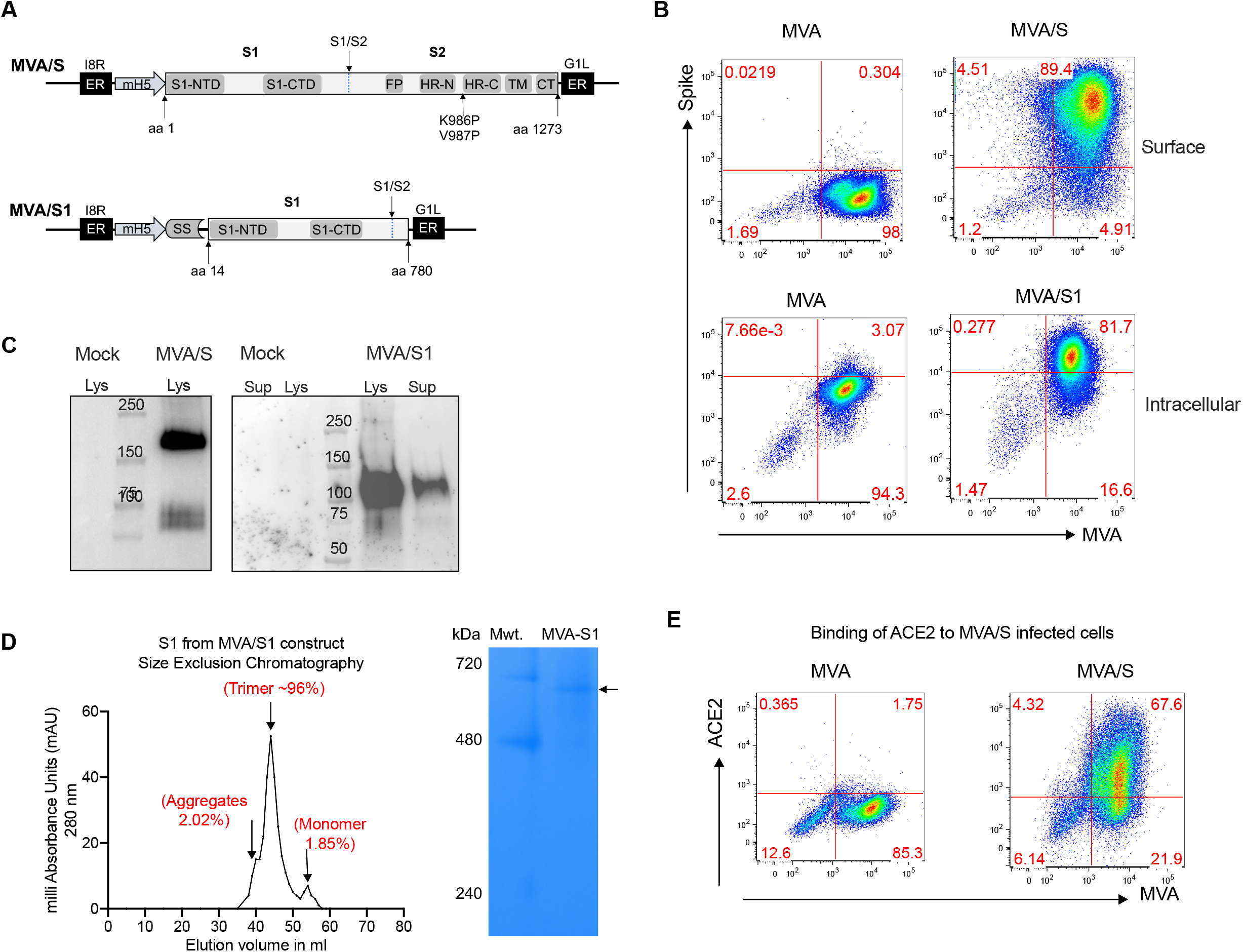
Construction and characterization of MVA/S and MVA/S1 recombinants. (A) Schematic representation of MVA/S and MVA/S1. Recombinant inserts were cloned in the essential region in between 18R and G1L under mH5 promoter. (B) Representative flow plots showing the expression of membrane anchored spike on the surface and S1 intracellularly. (C) Westernblotting analysis of expressed proteins in supernatants and lysates of MVA infected cells. (D) Size-exclusion chromatography analysis of S1 protein expressed by MVA/S1. (E) Binding of hACE2 to MVA/S expressing cells.

### Both MVA/S and MVA/S1 vaccines induce a strong binding antibody response but with different specificities

We immunized Balb/c mice with MVA/S or MVA/S1 on weeks 0 and 4, and measured binding antibody responses to total and different parts of spike i.e. RBD, S1, and S (S) using ELISA at 2 weeks post prime and boost (**Fig. 2**). While both vaccines induced a strong binding antibody response to S, they differentially targeted binding to RBD and S1 (**Fig. 2A, 2B**). The MVA/S sera showed binding to RBD whereas MVA/S1 sera showed binding to S1. This was interesting considering that S1 protein includes the RBD region and suggests that antibody binding in sera from MVA/S1-vaccinated mice may be targeting regions outside of the RBD. We also performed Luminex assay using sera obtained 3 weeks post-boost to measure binding to different parts of S including S2, and to determine the antibody subclass and their ability to bind different soluble FcγRs (**Fig. 2C**). These analyses revealed that MVA/S-vaccinated mice generated an antibody response that showed similar binding to the RBD, S1 and S2 purified proteins whereas serum from MVA/S1-vaccinated mice primarily generated an antibody response that bound to the non-RBD portion of the S1 (**Fig. 2C**). While the lack of binding to S2 is expected, poor binding to RBD was not expected. Analysis of IgG subclass and FcγR binding of RBD-specific antibody showed strong IgG2a response (Th1 biased) and binding to all three FcγRs tested with strongest binding to FcγR2 and FcγR4 in the MVA/S group (**Fig. 2D**). In contrast, poor binding of RBD-specific antibody was observed in general with MVA/S1 sera. However, the S1-specific antibody showed similar results in both groups. These results demonstrated differential targeting of spike specific antibody with Th1 profile induced by MVA/S and MVA/S1 vaccines.

**Fig 2.**
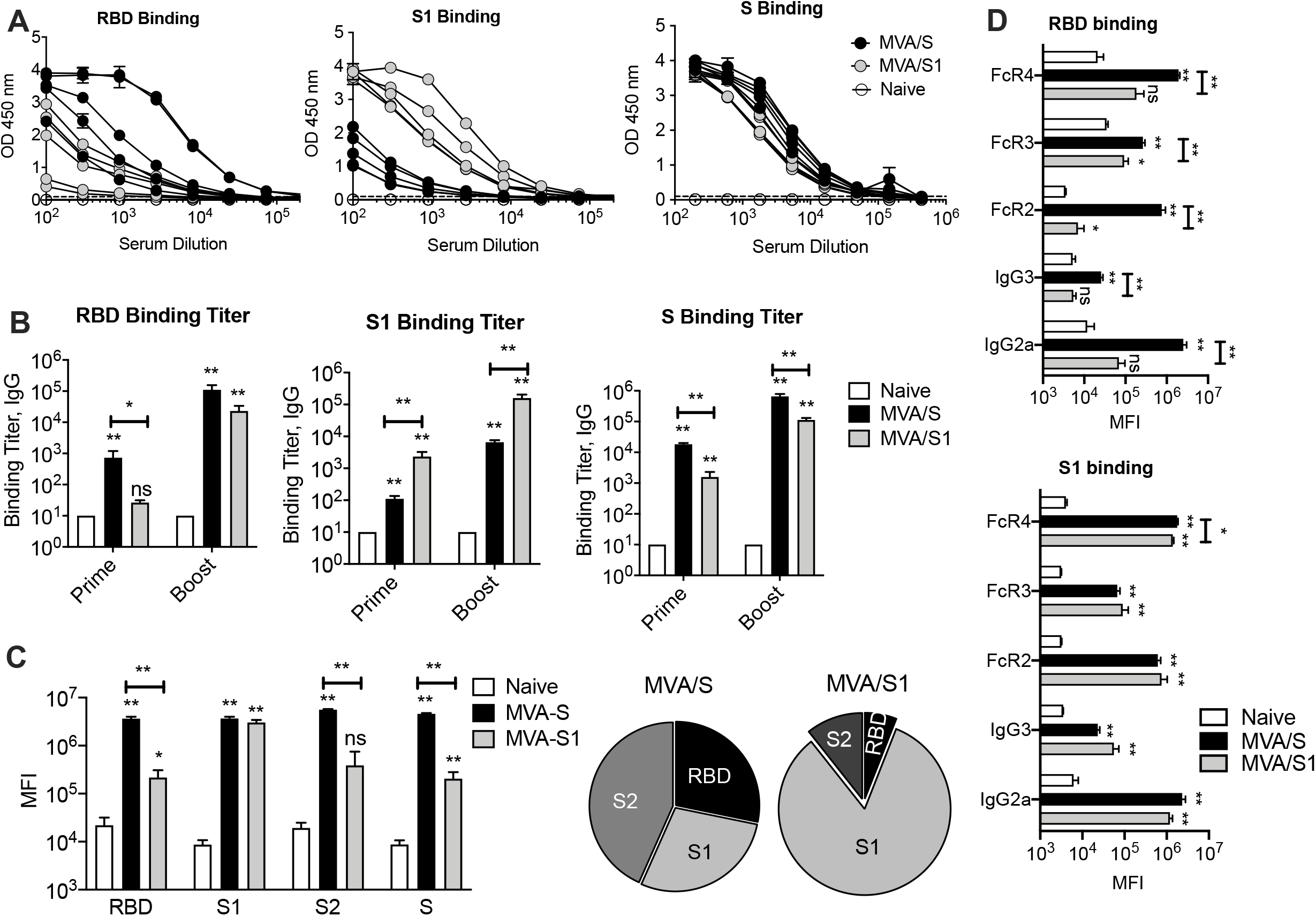
Antibody responses induced by MVA/S or MVA/S1 in mice. BALB/c mice were immunized on week 0 and 3 with recombinant MVAs expressing either S (MVA/S) (n=5) or S1 (MVA/S1) (n=5) in a prime-boost strategy. Unvaccinated (naïve) animals served as controls (n=5). (A) Binding IgG antibody response for individual proteins measured using ELISA at two weeks after boost. (B) Endpoint IgG titers against SARS-CoV-2 RBD, S1 and S measured at week 2 after immunization. The data show mean response in each group (n = 5) ± SEM. (C) Binding antibody response determined using Luminex assay at 3 weeks post boost. The pie graphs show the relative proportions of binding to three proteins in each group. (D) IgG subclass and soluble Fc receptor binding analysis of RBD and S1 specific IgG measured using the Luminex assay. Raw values are presented as in mean fluorescence intensity (MFI) in bar graph. The data represent mean responses in each group (n = 5) ± SEM.

### MVA vaccination induces strong bronchus-associated lymphoid tissue and antibody responses in the lung

We next studied if vaccination induced immune responses in the lung, a primary site of SARS-CoV-2 virus exposure. We measured the formation of bronchus-associated lymphoid tissue (BALT)*(22, 23)* using the immunohistochemistry at 3 weeks after the MVA boost by staining for B and T cells (**Fig. 3A, 3B**). As expected, the naïve mice showed very little or no BALT, however, the MVA vaccinated mice showed significant induction of BALT indicating the generation of local lymphoid tissue. While we do not know the longevity of persistence of these BALT, they are expected to help with rapid expansion of immunity in the lung following exposure to SARS-CoV-2 infection *(22, 24)*. Consistent with BALT, we also observed the induction of spike specific IgG responses in the BAL (**Fig. 3C**). Similar to serum, the RBD-specific IgG responses were higher in the MVA/S group compared to MVA/S1 group. These results demonstrate strong induction of antibody responses in the lung following MVA vaccination.

**Fig 3.**
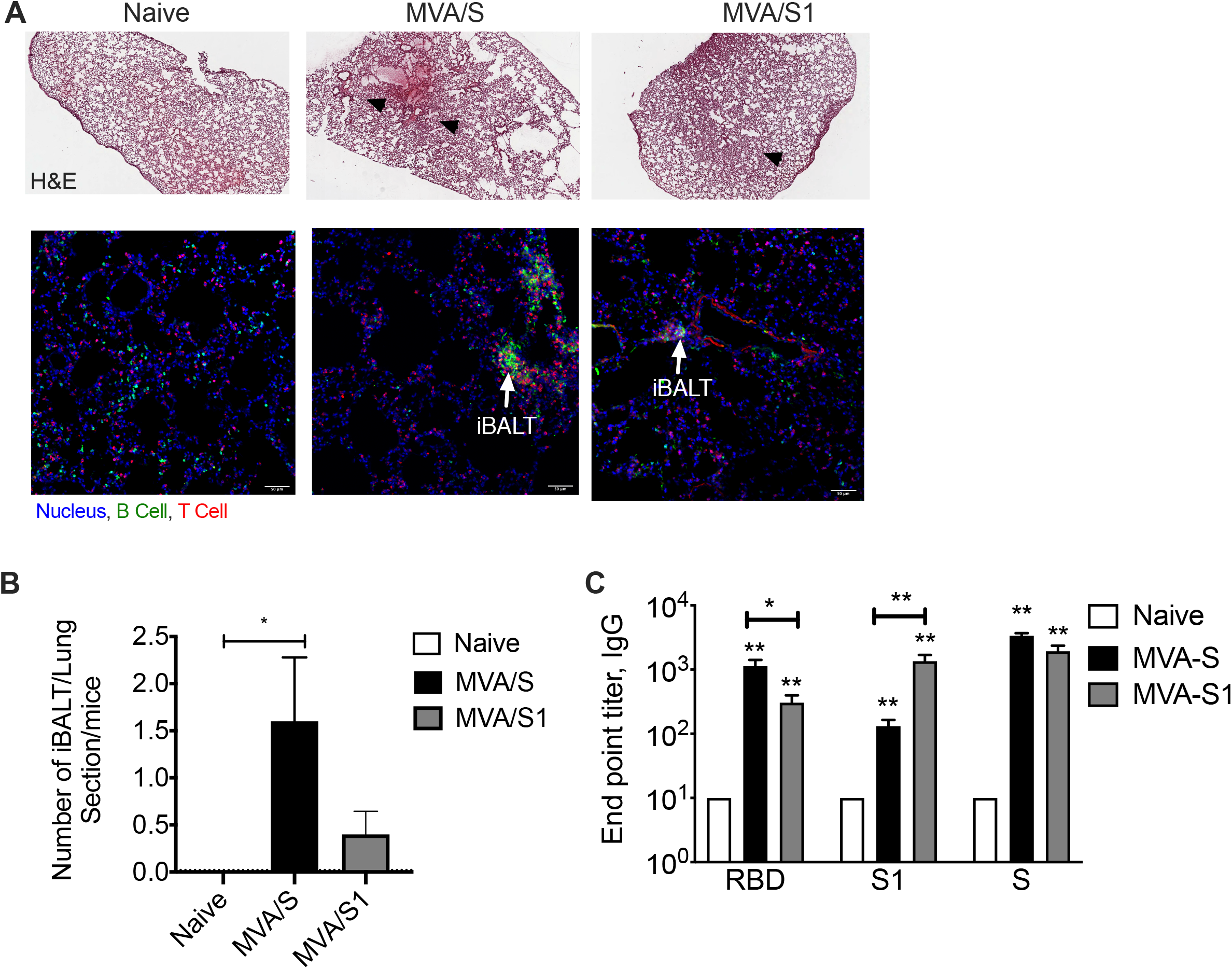
Inducible Bronchus Associated Lymphoid Tissues (iBALT) formation upon MVA/S and MVA/S1 vaccination. Frozen lung sections from vaccinated mice were either stained for H&E to analyze tissue structure and formation of iBALT aggregates (A), or immunofluorescence stained to visualize B cell and T cell (B) forming B cell follicle like structure (iBALT) induced by MVA/S vaccination given via i.m. route (right panel), and compared with unvaccinated control mice (left panel). Total number of iBALT like structures visualized in each section per mice was quantified and compared between the groups (C). The p value was calculated using non parametric mann-whitney test. (D) Lung immune responses in bronchoalveolar lavage (BAL) samples collected after euthanizations (three weeks post-boost) were measured using ELISA. SARS-CoV-2 S protein-specific binding IgG and IgA antibodies measured, and titters were presented in column graphs. The data represent mean responses in each group (n = 5) ± SEM.

### MVA/S but not MVA/S1 induces strong neutralizing antibody response

We next evaluated the neutralization capacity of serum from MVA-vaccinated mice using SARS-CoV-2. Using serum from 2 weeks post-boost (**Fig. 4A**), we performed an FRNT-mNG assay. Impressively, we observed a robust neutralizing antibody response in sera from mice vaccinated with MVA/S that ranged from 20-900 with a median of 200 (**Fig. 4B**). In contrast, we did not observe any detectable neutralization activity in sera from mice immunized with MVA/S1. This was despite the fact that MVA/S1 mice showed comparable or higher binding antibody response to RBD, S1 and S proteins. The neutralization titer correlated directly with the RBD binding titer and negatively correlated with the S1 binding titer (**Fig. 4C**). In particular, they correlated with the RBD or S specific IgG2a binding titer (**Fig. 4D**). These results demonstrated that MVA/S immunogen can induce a strong neutralizing antibody response against SARS-CoV-2 and could serve as a potential vaccine for SARS-CoV-2. Importantly, they also reveal that MVA/S1 is not a good vaccine as it failed to induce a neutralizing antibody response.

**Fig 4.**
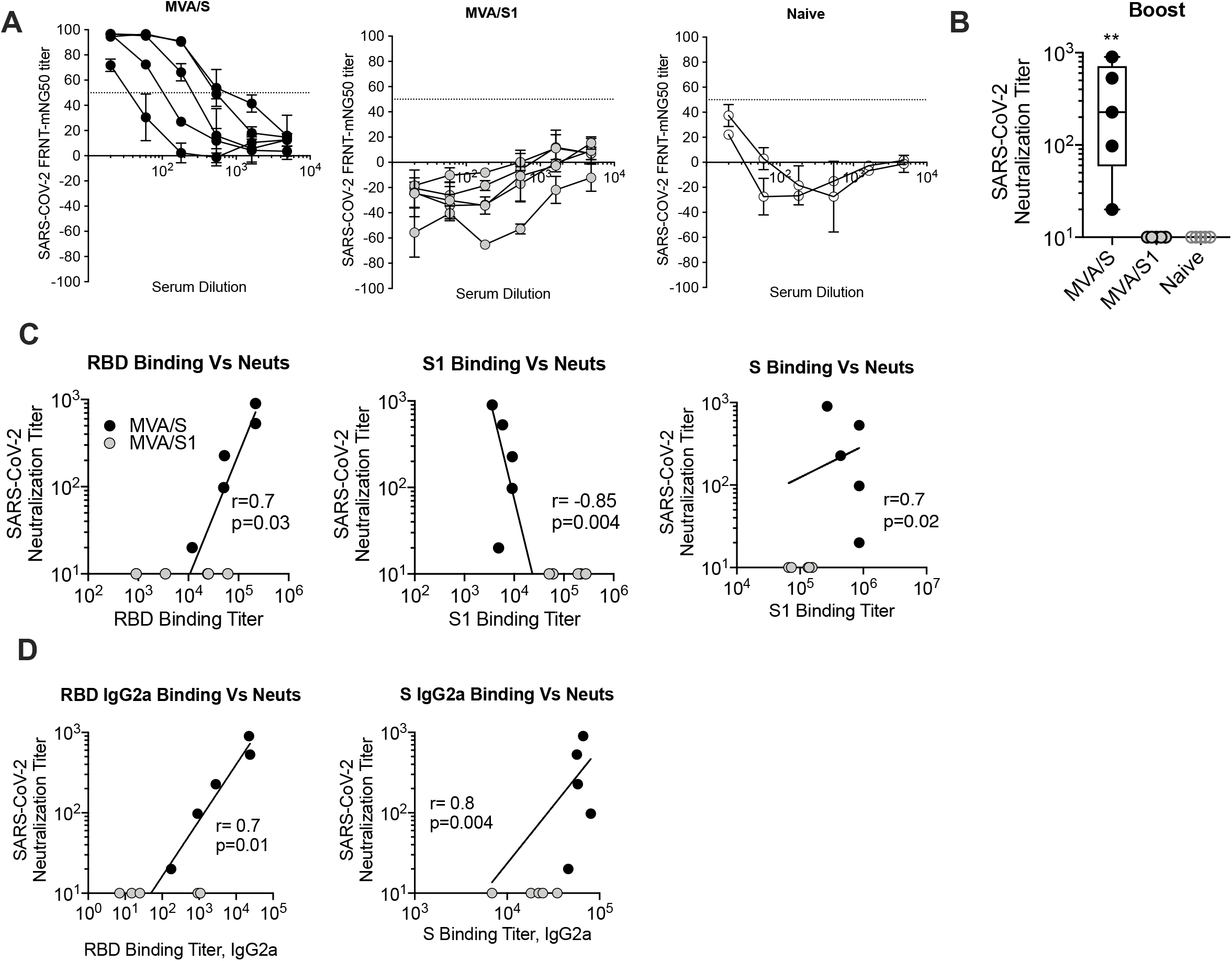
Neutralizing activity against SARS-CoV-2. (A) Percent neutralization of SARS-CoV-2 virus expressing GFP. Serum collected from the naïve animals used as negative controls. (B) Neutralization titer against SARS-CoV-2 virus expressing GFP. (C, D) Correlations between neutralization titer and ELISA binding titer.

### SARS-CoV-2 S1 exhibits lower affinity to ACE2 than RBD, which further weakens upon preincubation at 25°C

To further understand the failure of MVA/S1 vaccine to induce strong RBD binding antibody and neutralizing antibody, we purified the S1 trimer protein expressed by MVA/S1 vaccine and determined its ability to bind to human ACE-2 using biolayer interferometry (BLI)(**Fig. 5**). We used purified RBD protein as a benchmark. SARS-CoV-2 S1 bound to hu-ACE2 quite strongly but at 2-fold lower affinity than RBD (K_D_ = 70.1nM and 36nM respectively, **Fig. 5A, 5B**). S1 exhibited 10-fold lower association rate than RBD (k_on_ 1.1E+04 and 1.3E+05 1/Ms respectively). However, the affinity of S1-ACE2 further decreased by 5-fold when the protein was incubated at 25°C for 60 min. In contrary, RBD was stable and retained its affinity (K_D_ = 24nM). The data indicated the receptor binding domain of S1 to be unstable, thereby loosing association with ACE2 protein upon prolonged incubation at room temperature, unlike RBD. We observed 10-fold reduction in the association rate for S1-ACE2, compared to RBD which was meagerly affected.

**Fig 5.**
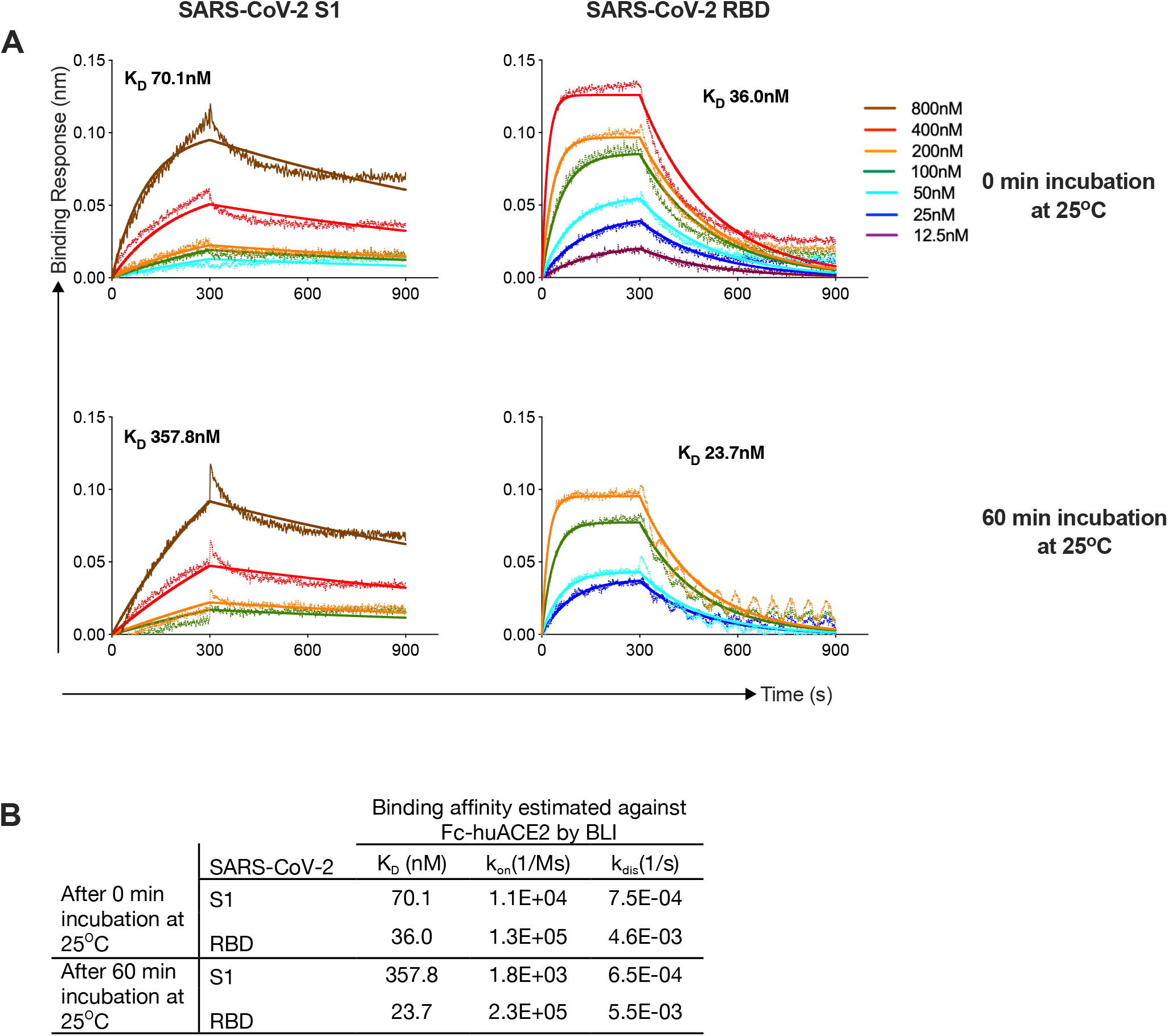
Analyzing SARS-CoV-2 RBD and S1 proteins affinities to human ACE2 (hACE2) proteins using biolayer interferometry (BLI). (A) Bio-Layer Interferometry sensograms of the binding of SARS-CoV-2 S1 and RBD proteins to immobilized Fc-human ACE2, after incubation of the analytes at 25°C for 0and 60 minutes. The traces represent BLI response curves for SARS-CoV-2 proteins serially diluted from 800nM to 12.5nM, as indicated. Dotted lines show raw response values, while bold solid lines show the fitted trace. Association and dissociation phases were monitored for 300s and 600s, respectively. The data was globally fit using a 1:1 binding model to estimate binding affinity. (B) Binding affinity specifications of S1 and RBD proteins against hu-ACE2.

## Discussion

Our results comparing two versions of MVA vaccines expressing either the full length prefusion stabilized spike or secreted S1 demonstrated that while both immunogens induce strong binding antibody response to spike only the former induces a strong neutralizing antibody response against the SARS-CoV-2. The failure of MVA/S1 immunogen to induce neutralizing activity was associated with its failure to induce antibody to RBD. This was surprising given the fact that RBD is part of S1. Binding to ACE-2 revealed that S1 presents RBD in the proper confirmation however the stability of RBD binding to its receptor seems to decrease markedly during prolonged incubation at the room temperature. This instability of S1 protein seems to contribute to induction of strong binding antibody to other regions in S1 other than RBD following immunization. Further studies are needed to understand the binding specificity of antibodies induced by these two vaccines. Importantly, systemic MVA vaccination also induced T cell and antibody responses in the lung that will be critical for protection against respiratory infections such as SARS-CoV-2. Our previous studies with MVA using HIV recombinants showed that MVA immunogenicity is durable and comparable in mice, non-human primates and humans except that we will need 10 times higher dose in NHPs and humans compared to mice *(16, 25)*. This accounts for a dose of about 10^8^ pfu and this dose is achievable by large scale manufacture. Collectively these results demonstrate that MVA/S could serve as a potential vaccine for SARS-CoV-2 and COVID-19.

## Methods

### Generation of recombinant MVAs expressing SARS-CoV 2 spike antigens

For MVA/S, the 3821-nt ORF encoding the SARS-nCoV Spike gene was codon optimized for vaccinia virus expression, synthesized (GenScript), and cloned into pLW-73 (provided by B. Moss and L. Wyatt, National Institutes of Health) using the XmaI and BamH1site under the control of the vaccinia virus modified H5 early late promoter and adjacent to the gene encoding enhanced GFP (green fluorescent protein) regulated by the vaccinia virus P11 late promoter. Similarly, to develop MVA/S1, spike secreted monomeric form, GmCSF signal sequence followed with Spike DNA sequence of 14-780 AA was synthesized and cloned between Xma1 and BamH1 sites of pLW-73 vector. These plasmid DNAs were subsequently used to generate recombinant MVAs by transfecting transfer plasmids into DF-1 cells that were infected with 0.05 plaque forming units (pfu) of MVA per cell as described previously into the essential region of MVA 1974 strain between genes I8R and G1L *(26)*. Recombinant MVAs (rMVA) were isolated using standard methods. The expression of spike on the MVA-infected cells from MVA/S construct was determined using anti-SARS-CoV-2 spike antibody (Cat #135356, GeneTex). For MVA/S1, anti SARS-CoV-2 RBD antibody (Cat# 40592-T62, Sino Biological) was used to stain intracellularly. Plaques were picked for 7 rounds to obtain GFP-negative recombinants and DNA was sequenced to confirm there are no mutations. Viral stocks were purified from lysates of infected DF-1 cells using a 36% sucrose cushion and titrated using DF-1 cells by counting pfu/ml. Absence of wildtype MVA was confirmed by PCR using recombinant specific primers flanking the inserts, and by flow cytometry by staining for MVA and spike.

### Flow staining for Spike expression

DF-1 cells were infected with MVA/S or MVA/S1 at 1 pfu per cell and cells were stained after 36h of infection. MVA/S infected cells were harvested, stained with live dead marker and anti SARS-CoV-2 spike antibody (#GTX135356, GeneTex) followed by donkey anti-rabbit IgG coupled to PE (# 406421, BioLegend) on the surface. Cells were then fixed with Cytofix/cytoperm (BD Pharmingen), permeabilized with permwash (BD Pharmingen), and intracellularly stained for MVA using mouse monoclonal anti-Vaccinia virus E3L Ab (#NR-4547 BEI Resources) coupled to PacBlue. MVA/S1 infected cells were stained for live dead marker, fixed, permeabilized and intracellularly stained for S1 protein using SARS-CoV-2 RBD Ab (#40592-T62, SinoBiological), followed by sequentially with donkey anti Rabbit PE and anti – Vaccinia virus E3L Ab. Spike and MVA double positive cells were shown on live cells. Similarly human ACE2 binding to surface expressed spike on MVA/S infected DF-1 cells was done using biotinylated human ACE2 protein (#10108-H08H-B, Sino Biological) followed by streptavidin-PE (BD Pharmingen) and intracellularly for MVA as described earlier.

### Protein expression and purification

RBD-His and S1 proteins were produced in Amara laboratory by transfecting HEK293 cells using plasmids pCAGGS-RBD-His and pGA8-S1, respectively. The RBD-His plasmid was obtained from BEI resources (Cat# NR-52309). The pGA8-S1 plasmid was generated by cloning human codon-optimized S1 DNA sequence from amino acids 14-780 with GM-CSF signal sequence under the control of CMV promoter with intron A. Transfections were performed according to manufacturer’s instructions (Thermo Fischer). Briefly, HEK 293F cells were seeded at a density of 2×10^6^ cells/ml in Expi293 expression medium and incubated in an orbital shaking incubator at 37°C and 127 rpm with 8% CO_2_ overnight. Next day, 2.5×10^6^ cells/ml were transfected using ExpiFectamine™ 293 transfection reagent (ThermoFisher, cat. no. A14524) as per manufacturer’s protocol. The cells were grown for 72hrs at 37°C,127 rpm, 8% CO_2_. The cells were harvested and spun at 2,000g for 10 minutes at 4°C. The supernatant was filtered using a 0.22 μm stericup filter (Themofischer, cat.no. 290-4520), and loaded onto pre-equilibrated affinity column for protein purification. The SARS-CoV-2 RBD-His tag and S1 proteins were purified using Ni-NTA resin (Thermofischer, cat.no. 88221) and Agarose bound Galanthus Nivalis Lectin (GNL) (Vector Labs, cat. no. AL-1243-5) respectively. Briefly, His-Pur Ni-NTA resin was washed with fresh PBS by centrifugation at 2000g for 10 min. The resin was resuspended with the supernatant and incubated for 2 hours on a shaker at RT. Polypropylene column was loaded with the supernatant-resin mixture and then washed with wash Buffer (25mM Imidazole, 6.7mM NaH_2_PO_4_.H_2_O and 300mM NaCl in PBS) four times, after which the protein was eluted in elution buffer (235mM Imidazole, 6.7mM NaH_2_PO_4_.H_2_O and 300mM NaCl in PBS). S1 protein supernatants were mixed with GNL-resin, overnight on rocker at 4°C. The supernatant-resin mix was loaded on to the column and washed 3 times with PBS and eluted using 1M methyl-α-D mannopyranoside, pH7.4. Protein elutions were dialysed against PBS using Slide-A-lyzer Dialysis Cassette (ThermoScientific, cat no. 66030), concentrated using 10 kDa Amicon Centrifugal Filter Units (for RBD) or 50 kDa Amicon Centrifugal Filter Units (for S1) at 2000g at 4°C. The concentrated protein elutes were run on a Superdex 200 Increase 10/300 GL (GE Healthcare) column on an Akta™Pure (GE Healthcare) system to collect the peaks of interest. The eluted peaks of interest were pooled. The concentration was estimated by BCA Protein Assay Kit (Pierce). The purity of the protein was confirmed by BN-PAGE (NuPAGE™, 4-12% Bis-Tris Protein Gels, ThermoScientific), SDS PAGE and Western blot.

### Western Blotting

DF-1 cells were infected with 1 MOI of recombinant MVA/S or MVA/S1 for 36 h. Infected cells were lysed in ice-cold RIPA buffer and supernatants were collected. Lysates were kept on ice for 10 min, centrifuged, and resolved by SDS PAGE using precast 4–15% SDS polyacrylamide gels (BioRad). Proteins were transferred to a nitrocellulose membrane, blocked with 1% casein blocker overnight (Cat#1610782 Biorad), and incubated for 1 h at room temperature with anti-SARS-CoV-2 spike mouse mAb (Cat # GTX632604, GeneTex) for MVA/S and rabbit SARS-CoV-2 RBD polyclonal antibody (Cat# 40592-T62, Sino Biological) for MVA/S1 diluted 1:2500 in blocking buffer, respectively. The membrane was washed in PBS containing Tween-20 (0.05%) and was incubated for 1 h with horseradish peroxidase-conjugated anti-mouse or anti-rabbit secondary antibody (Southern Biotech) diluted 1:20,000 accordingly. The membranes were washed, and proteins were visualized using the ECL select chemiluminescence substrate (Cat# RPN2235 GEhealthcare).

### Animal immunizations and sample collection

BALB/c mice were obtained from Jackson Laboratories (Willmington, MA, USA) and were housed in the animal facility at the YNPRC of Emory University, Atlanta, GA. All animal experiments were reviewed and approved by the Animal Care and Use Committee of the Emory University. Briefly, 6–8-week-old female BALB/c mice (n=5) were immunized with 10^7 plaque-forming-units (pfu) of rMVA/S or rMVA/S1 by intramuscular (i.m.) route on weeks 0 and 3. Blood samples were collected by facial vein puncture in BD Microtainer^®^ Tube at two weeks after each immunization for analyzing serum antibody responses. At three weeks after the boost, animals were euthanized using CO_2_ followed by cervical dislocation. Blood, lung tissue and bronchoalveolar lavage (BAL) were collected. A small piece of lung was used for IHC staining.

### Measurement of binding antibodies by enzyme-linked immunosorbent assay (ELISA)

These were performed as described previously *(27)*. Briefly, Nunc high-binding ELISA plates (Thermo Fisher Scientific; Waltham, MA, USA) were coated with 2 μg/ml of recombinant SARS-CoV-2 proteins (S1 and RBD-His proteins were produced in the lab and S1 +S2 ECD spike protein was purchased from Sino Biologicals) in Dulbecco’s phosphate-buffered saline (DPBS) and incubated overnight at 4 °C. Plates were then blocked with 5% blotting-grade milk powder (Sigma-Aldrich; St. Louis, MO) and 4% whey powder (Sigma-Aldrich; St. Louis, MO) in DPBS with 0.05% Tween 20 for 2 h at room temperature (RT). Plates were then incubated with serial dilutions of mouse sera for 2h at RT, washed 6 times and then incubated with 1: 6000 dilution of horseradish peroxidase (HRP) conjugated anti-mouse IgG secondary antibody (Southern Biotech; Birmingham, AL, USA) for 1 h at RT. The plates were washed and developed using TMB (2-Component Microwell Peroxidase Substrate Kit, SeraCare’s Reagents; Milford, MA, USA) and the reaction was stopped using 1N Sulfuric acid solution. Plates were read at 450 nm wavelength within 30 min using ELISA plate reader (Molecular Devices; San Jose, CA, USA). ELISA endpoint titers were defined as the highest reciprocal serum dilution that yielded an absorbance >2-fold over background values.

### Measurement of SARS-CoV-2 neutralizing antibodies

Infectious clone of the full-length mNeonGreen SARS-CoV-2 (2019-nCoV/USA_WA1/2020) was generated as previously described *(28)*. The stock virus was passaged twice (P2) and viral titers were determined by plaque assay on Vero E6 cells (ATCC). Vero cells were cultured in complete DMEM medium consisting of 1x DMEM (Corning Cellgro), 10% FBS, 25mM HEPES Buffer (Corning Cellgro), 2mM L-glutamine, 1mM sodium pyruvate, 1x Non-essential Amino Acids, and 1x antibiotics. Mouse sera from vaccinated animals and naive animals were heat inactivated for 30 min at 56 °C. Heat-inactivated serum was serially diluted in duplicate three-fold starting at a 1:20 dilution in a 96-well round-bottom plate and incubated between 400-600 FFU of ic-SARS-CoV-2-mNG for 1 h at 37°C. This antibody-virus mixture was transferred into the wells of a 96-well plate that had been seeded the previous day at 2.5× 10^4^ cells per well. After 1 hour, the antibody-virus inoculum was removed and 0.85% methylcellulose in 2% FBS containing cDMEM was overlaid onto the cell monolayer. Cells were incubated at 37°C for 24-28 hours. Cells were washed three times with 1x PBS (Corning Cellgro) and fixed with 125 μl of 2% paraformaldehyde (Electron Microscopy Sciences) for 30 minutes. Following fixation, plates were washed twice with 1x PBS and imaged on an ELISPOT reader (CTL Analyzer). Foci were counted using Viridot (counted first under the green light setting followed by background subtraction under the red light setting) *(29)*. FRNT50 curves were generated by non-linear regression analysis using the 4PL sigmoidal dose curve equation on Prism 8 (Graphpad Software). Neutralization titers were calculated as 100% x [1-(average foci in duplicate wells incubated with mouse serum) ÷ (average number of foci in the duplicate wells incubated at the highest dilution of the respective mouse serum)].

### Immunohistochemistry and Confocal microscopy

Tissue imaging of optimum cutting temperature (OCT) compound embedded lung tissue sections was performed. Fresh lung tissues from vaccinated and control mice were fixed in 4% PFA for 60 minutes at RT followed by overnight 30% sucrose treatment. Next day, the tissues are embedded in OCT compound and and snap frozen. 10-micron thick tissue sections were taken from Frozen OCT blocks and adhered to highly adhesive glass slides. Tissue sections were immediately fixed in 30% PFA and 70% acetone fixative cocktail. Fixed tissue sections were subjected to blocking with 5% BSA in PBS supplemented with 2% goat serum and 0.025% Tritox-100 for 1h at RT. After three washes with chilled PBS, tissue sections were subjected to overnight incubation with primary antibodies in 1% BSA and 0.025% Triton-100 supplemented PBS. For investigating the formation of iBALT B cell and T cells were stained in lungs using primary antibody cocktail containing rat anti-mouse CD45R/B220 (BioLegend, Cat#103202) and Hamster anti mouse CD3 (BD, Cat#550277) antibodies. and incubated overnight at 4 degree Celsius. Next day, primary antibodies were washed off the slides thrice using chilled PBS followed by incubation with secondary antibodies for 1h at RT. Secondary antibodies cocktail contained goat antirat IgG-Alexa488 (abcam, Cat#ab150157) and goat anti-hamster IgG-Alexa546 (Thermofisher, Cat#A21111). Slides were washed thrice with chilled PBS and mounted using anti-fade mounting media containing DAPI to stain nucleus. The slides were imaged by Olympus FV1000 confocal imaging using 20X objective and images were analyzed using ImageJ.

### Systems serology by Luminex analysis

For relative quantification of antigen-specific antibody titers, a customized multiplexed approach was applied, as previously described *(30)*. Therefore, magnetic microspheres with a unique fluorescent signature (Luminex) were coupled with SARS-CoV-2 antigens including spike protein (S) (provided by Eric Fischer, Dana Farber), Receptor Binding Domain (RBD), and CoV HKU1 RBD (provided by Aaron Schmidt, Ragon Institute), CoV-2 S1 and S2 (Sino Biologicals) as well as influenza as control (Immune Tech). Coupling was performed using EDC (Thermo Scientific) and Sulfo-NHS (Thermo Scientific) to covalently couple antigens to beads. 1.2×10^3^ beads per region/ antigen were added to a 384-well plate (Greiner), and incubated with diluted plasma samples (1:90 for all readouts) for 16h shaking at 900rmp at 4°C to facilitate immune complex formation. The next day, immune complexed microspheres were washed three times in 0.1% BSA and 0.05% Tween-20 using an automated magnetic plate washer (Tecan). Anti-mouse IgG-, IgG2a-, IgG3-, IgA- and IgM-PE coupled (Southern Biotech) detection antibodies were diluted in Luminex assay buffer to 0.65ug/ml. Beads and detection antibody were incubated for 1h at RT while shaking at 900rpm. Following washing of stained immune complexes, a tertiary goat anti-mouse IgG-PE antibody (Southern Biotech) was added and incubated for 1h at RT on a shaker. To asses Fc-receptor binding, mouse Fc-receptor FcγR2, FcγR3, FcγR4 (Duke Protein Production facility) were biotinylated (Thermo Scientific) and conjugated to Streptavidin-PE for 10 min (Southern Biotech) before adding to immune complexes. Finally, beads were washed and acquired on a flow cytometer, iQue (Intellicyt) with a robot arm (PAA). Events were gated on each bead region, median fluorescence of PE for bead positive events was reported. Samples were run in duplicate for each secondary detection agent.

### Binding kinetics by Bio-layer Interferometry (BLI)

The assay was done in 384 well format on Octet Red384 platform, Pall ForteBio. Fc-human ACE2 (Acro Biosystems, catalogue no AC2-H5257) was immobilized onto anti-human Fc coated biosensors at 5μg/ml. The purified SARS-CoV-2 protein (S1, RBD) was used as analyte (800nM-12.5nM). The analyte was incubated at 25°C for either 0min and 60min before running the experiment. Protein-ACE2 association (k_on_) was monitored for 300s, followed by dissociation (k_dis_) for 600s. 10X Kinetics Buffer, Pall ForteBio was used as buffer for all steps and dilutions in the assay. K_D_ (dissociation constant = k_dis_/k_on_) was estimated by globally fitting the reference (buffer only) subtracted sensograms to a 1:1 binding model using ForteBio Data analysis v9 software.

### Data analysis and statistical procedures

GraphPad Prism v7.0 was used to perform data analysis and statistics. A Mann-Whitney rank sum test was used to compare between Naïve, MVA/S and MVA/S1 vaccination groups. P value of <0.05 was considered significant. The correlation analysis was performed using Spearman rank test.

## Acknowledgements

We thank Drs. Bernard Moss and Lynda Wyatt for providing the pLW73 transfer plasmid, the Yerkes Division of Research Resources for animal care and Histology and Molecular Pathology Lab for help with tissue sectioning. The following reagent was produced under HHSN272201400008C and obtained through BEI Resources, NIAID, NIH: Vector pCAGGS Containing the SARS-Related Coronavirus 2, Wuhan-Hu-1 Spike Glycoprotein Receptor Binding Domain (RBD), NR-52309.

## Funding

This work was supported in part by National Institutes of Health Grants RO1 AI148378-01S1 to R.R.A., and NCRR/NIH base grant P51 OD011132 to YNPRC.

## Author Contributions

R.R.A. was responsible for overall experimental design and supervision of laboratory studies, manuscript writing and editing. N.R, S.G, N.C, A.S, S.A.R, A.S, S. F, and C.A were responsible for conducting experiments, data collection and data analysis. G.A supervised Luminex assays. M.S conducted and supervised neutralizing antibody assays. All authors contributed to manuscript writing and editing.

## Notes

### Competing Interest Statement

The authors have declared no competing interest.

